# LTP-patterned electromagnetic stimulation induces NMDA receptor-dependent synaptic plasticity in cortical networks

**DOI:** 10.64898/2026.06.17.732958

**Authors:** Cooper Kansala, Jesse St. Jean, Vanessa Nkansah-Okoree, Nicolas Rouleau, Nirosha J. Murugan

## Abstract

Bioinspired electromagnetic stimulation, in which fields are patterned after endogenous neural activity, have emerged as a potential non-invasive approach for modulating brain dynamics, yet the waveform parameters that determine biological specificity remain poorly defined. Complex electromagnetic patterns modeled after long-term potentiation (LTP) have been reported to alter learning and cortical injury outcomes *in vivo*, yet whether these fields engage cell-scale synaptic plasticity mechanisms remains unclear. Here, we show that a microtesla-strength electromagnetic field (EMF) patterned after electrophysiological signatures of long-term potentiation produces complex waveform-specific changes in primary cortical network dynamics. Using high-density microelectrode arrays, we show that LTP-patterned EMF stimulation transiently increases spontaneous spikes-per-burst activity relative to a frequency-matched sine-wave EMF exposure and a sham, no field control. This waveform-dependent effect was abolished by NMDA receptor antagonism, indicating dependence on glutamatergic signaling pathways linked to activity-dependent plasticity. LTP-EMF stimulation also dynamically altered evoked network responses, reducing active electrode recruitment to direct electrical stimulation immediately after exposure, with recovery at later timepoints, consistent with a reversible post-induction reorganisation of network state. Transcriptional profiling identified a delayed adaptive response enriched for cellular remodeling pathways, and immunocytochemistry revealed increased co-localization of pre- and post-synaptic markers synaptophysin and PSD-95, a microstructural hallmark of synaptogenesis. Overall, these findings show that weak, bioinspired EMF stimulation can induce waveform-specific changes in cortical network dynamics and engage NMDA-dependent, plasticity-associated mechanisms. This work supports the perspective that temporal waveform structure is a key stimulation parameter for optimizing non-invasive electromagnetic modulation of neural activity.

## 1.0 Introduction

Non-invasive neuromodulation techniques, including pulsed electromagnetic fields (EMFs), have emerged as powerful tools to probe and alter brain function, with applications spanning cognitive research to the treatment of neuropsychiatric disease [1]. Yet clinical and experimental responses remain variable, even when stimulation is delivered using standardized protocols [2–4]. This variability is usually attributed to anatomical differences [5] or imprecise spatial targeting [6]. However, the temporal structure of the signal itself has not received similar consideration. Because neural networks encode and process information through temporally patterned activity [7], externally applied EMFs may be best understood as both energetic perturbations defined by intensity and frequency, as well as patterned, information-bearing signals capable of interacting with endogenous neural dynamics.

Brain tissues generate endogenous, time-varying EMFs that may inform the development of more biocompatible neurostimulation technologies. Indeed, coherent neuronal oscillations generate bulk electric fields of approximately 2 mV mm⁻¹ [8] and magnetic fields in the range of 25–100 nT [9, 10], detectable at the scalp as electro- and magneto-encephalographic signals. At the cellular scale, neurons produce local field potentials that depolarize adjacent membranes [11–15] and modulate firing in neighbouring cells through ephaptic coupling [16], synchronizing neurons and entraining oscillations across cortical and hippocampal circuits [17] with functional consequences for information processing and memory [18]. Endogenous fields as weak as 0.5 mV mm⁻¹ [19] can shift membrane potential by 0.1–1.3 mV [11, 20] and bias action potential timing. Thus, low-intensity EMFs within the range of endogenous fields may be sufficient to stimulate brain function when signal patterns reflect the temporal structure of physiological rhythms.

Neurons exist within dynamic biophysical tissue microenvironments, where endogenous time-varying EMFs are physiologically relevant and functionally consequential. One central feature of endogenous neural signaling is that information is encoded and processed as temporally patterned activity. Bursts, nested rhythms, and other complex waveforms act as physiological “control signals” that gate synaptic integration and plasticity [18]. Whether a synapse is strengthened or weakened depends on the precise timing of pre- and post-synaptic activity rather than on mean firing rate alone [20]. It may be hypothesized that temporal patterning is a relevant parameter of EMF-based neurostimulation that can be tuned to maximize effects.

Long-term potentiation (LTP) is a well-characterized example of temporal patterning as an instructive signal for plasticity within the brain [19]. Stimulation protocols that mimic endogenous burst activity, including theta-burst, and primed-burst patterns, lower the threshold for LTP induction and engage plasticity mechanisms that are not recruited equivalently by simple periodic stimulation trains [20]. Complex spike bursts and theta-patterned inputs, generate post-synaptic conditions that promote N-methyl-D-aspartate (NMDA) receptor activation and synaptic potentiation, even when mean firing rates remain modest [21] because this frequency is thought to preferentially disable feed-forward inhibition and promote post-synaptic depolarization [21]. These findings support the hypothesis that temporal patterning is biologically instructive and motivate its use as a meaningful parameter in applied neurostimulation design.

Transcranial magnetic stimulation (TMS) is a non-invasive method for modulating cortical excitability and has become an established high-intensity (>1 tesla) clinical tool for the treatment of select neuropsychiatric disorders. Repetitive low-intensity (<100 millitesla) magnetic stimulation induces Ca²⁺-dependent cellular responses in cortical slices and neuronal cultures [22, 23] and transcranial µT-strength magnetic fields were recently shown to suppress neuroinflammation and oxidative stress *in vitro* [24]. These findings support the principle that field intensity alone does not determine neurobiological efficacy, and that temporal structure is a separable design parameter. Response heterogeneity nevertheless remains a critical barrier for translation, suggesting that efficacy depends on whether the imposed stimulus couples meaningfully to endogenous signalling dynamics rather than on dosage alone [22, 25]. Biomimetic stimulation patterns, which recapitulate the complex bioelectric oscillations of neural tissue, represent an underexplored approach to addressing this problem [19, 26]. Early studies reported that EMFs patterned to replicate the bioelectric signatures of LTP impaired memory acquisition in rodents [27, 28] and attenuated excitotoxic cortical damage following lithium-pilocarpine-induced seizures [29]. However, whether such fields directly engage the cell-scale mechanisms of synaptic plasticity they were designed to recapitulate had not been investigated.

Here, we address this knowledge gap by testing the hypothesis that biomimetic field patterning, designed to match the endogenous plasticity-relevant temporal structure of LTP signaling, can selectively engage synaptic mechanisms of cortical cells *in vitro* more than frequency-matched but simple (i.e., low information) waveforms (**Fig. 1**). We compared a µT-intensity EMF patterned with LTP-associated waveform signatures, a 100 Hz sinewave exposure, and No Field (sham) controls (**Fig. 2**). Using primary embryonic rat cortical networks cultured on high-density microelectrode arrays (HD-MEAs), we assessed spontaneous bursting, evoked responses, NMDA receptor dependences, transcriptional responses, and synaptic marker organization. Our findings suggested that exposure to an LTP-patterned EMF increased spikes-per-burst within the networks while transiently suppressing excitability to applied voltages, which was abolished by NMDA receptor antagonists. These networks also displayed increased synaptophysin and PSD-95 co-localization relative to controls, suggestive of new synapse formation. Collectively, these findings support a stimulation design framework in which applied EMFs are treated as patterned signals rather than generic energy delivery systems. Consistent with this view, we suggest that biomimetic temporal structure is a controllable parameter that can gate synaptic and network-level outcomes, offering a new roadmap to targeted neuromodulation and a mechanistic foundation for clinical innovation.

**Figure 1.**
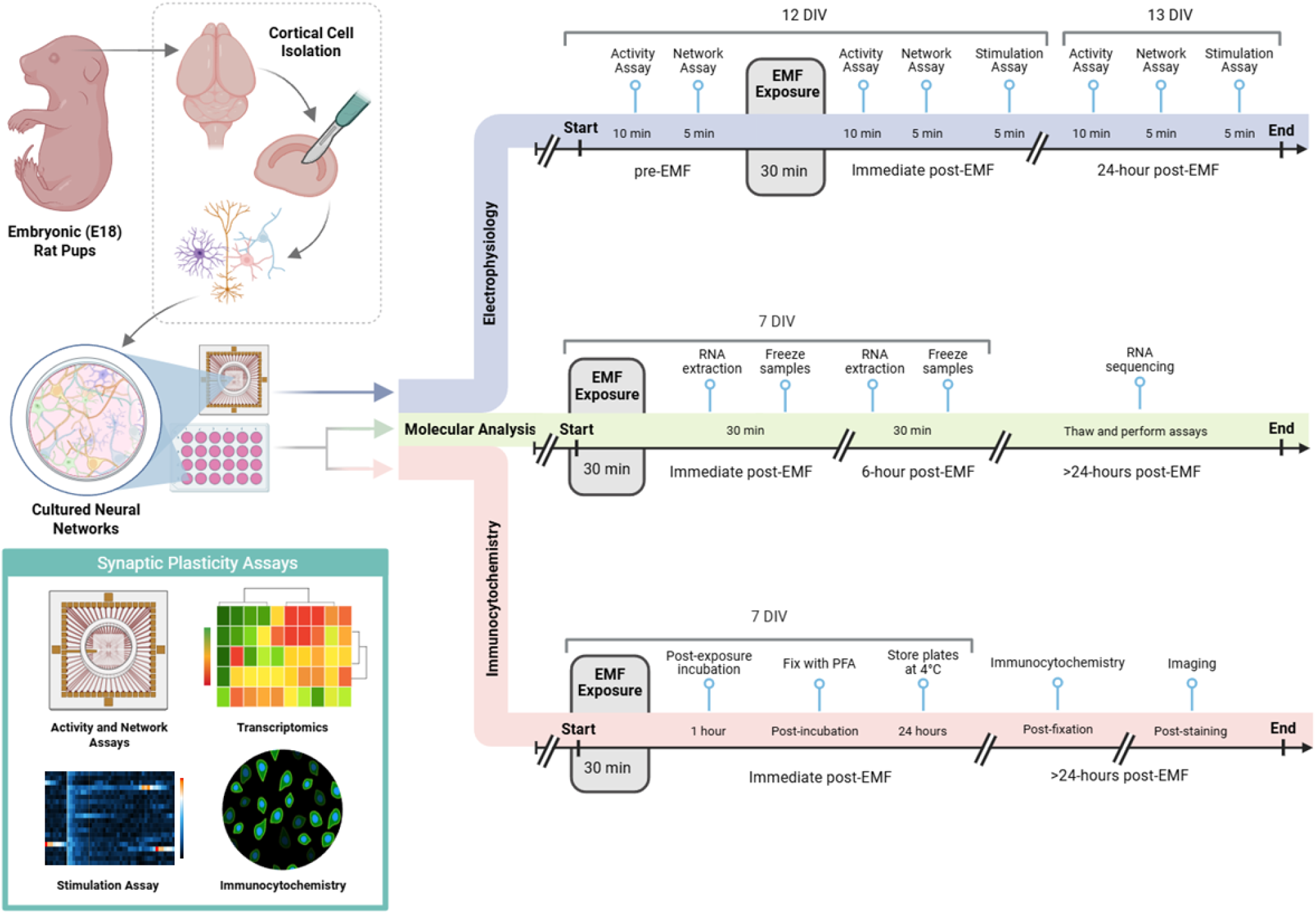
Experimental workflow to assess biomimetic EMF stimulation effects in primary neural cultures. Primary cortical cells, including neurons and glial cells, were isolated from E18 Sprague-Dawley rat embryos, seeded in 24-, 48-well plates or MaxOne HD-MEA chips, prior to patterned EM exposures. Post-exposure assays included 3 parallel pipelines: Electrophysiological recordings (blue path: Activity, Network, and Stimulation Assays), transcriptomic profiling (green path: RNAseq), and immunocytochemistry to visualize synapse formation (red path). As indicated by the workflow diagrams, assays were performed at various timepoints (7, 12, or 13 days *in vitro* or “DIV”). Oblique, double dashed lines indicate discontinuities for distinct phases of the workflow and may be of variable durations (e.g., frozen samples are tested days or weeks later).

**Figure 2.**
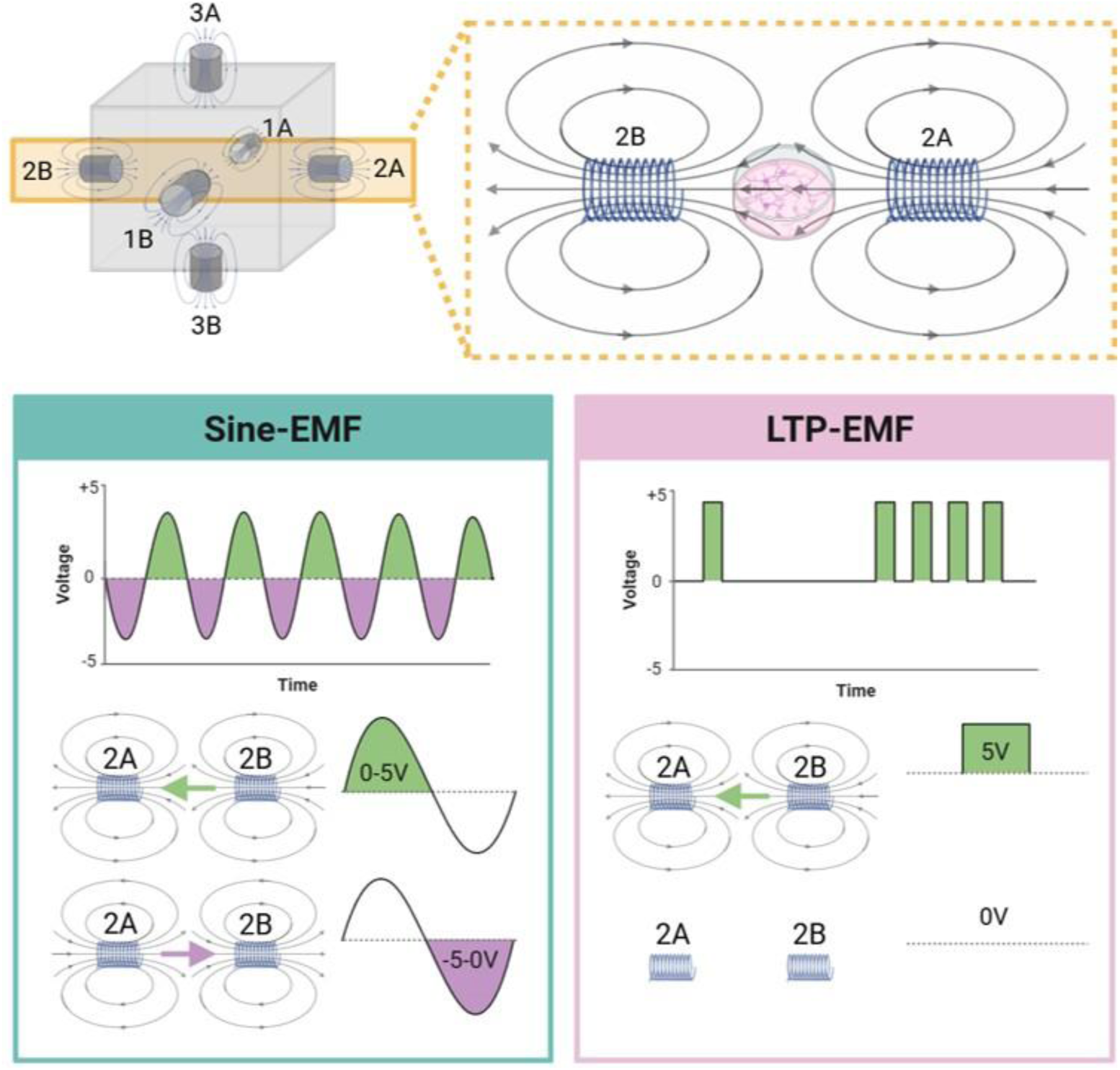
EMF exposure chamber and stimulation waveform configurations. Three orthogonally convergent solenoid pairs (1A-B, 2A-B, 3A-B), each positioned at 180° to one another, delivered weak µT-strength EMFs to a central focal point within the exposure chamber (top panel). All experiments were performed within an incubator (37°C, 5% CO_2_, 95% humidity). Cultured neural networks were exposed to a continuous 100 Hz sine wave (Sine-EMF, bottom left panel) or a complex pulse pattern modeled after the electrophysiological signatures of LTP (LTP-EMF, bottom right panel). While the Sine-EMF condition was associated with graded voltages between +5 and -5 V, including periodic reversal of the field direction, the LTP-EMF was monophasic, with pulses presented as all-or-none +5 V potentials, with interstimulus periods defined as 0 V output, corresponding to no-field events between pulses. A detailed description of the LTP-EMF waveform’s temporal parameters and endogenous analogs are described elsewhere. Unexposed controls (No Field) were placed within the exposure box for identical periods of time without activating the solenoids as a sham procedure.

## 2.0 Results

### 2.1 LTP-EMF exposures transiently increased spikes-per-burst in cultured neural networks

Previous *in vivo* studies reported that whole-body exposure to LTP-patterned electromagnetic fields impaired memory acquisition and reduced excitotoxic cortical damage in experimentally seized rodents [27–29], suggesting that temporally-patterned EMFs may engage with neural pathways tied to excitability and plasticity, yet these claims remained uncharacterized. To address this, we exposed primary rat cortical cultures grown on HD-MEAs to one of three conditions: an LTP-patterned EMF, a continuous 100 Hz sine-wave EMF, or a No Field sham exposure condition (**Fig. 2**) and performed spontaneous activity recordings and network burst analysis immediately, 6-hrs, and 24-hrs post-exposure. Each chip was measured at all time points as a within-subject, repeated measure that allowed standardization to baseline activity, accounting for chip-to-chip variability. LTP-EMF exposure produced a significant field-dependent effect on normalized spikes-per-burst across post-exposure time points, F(2,12)=5.76, p<0.05 (**Fig. 3A**). Additional analyses revealed the effect was driven by the immediate timepoint, H(2)=9.38, p<0.01. “Immediate”, in this context, refers to MEA measurements performed immediately upon completing the 30-min EMF exposure (or sham) and moving the chip from an exposure incubator to a measurement incubator within the same room. Post-hoc comparisons within post-exposure timepoints revealed that networks displayed more spikes-per-burst immediately after the LTP-EMF exposure relative to Sine-EMF (p<0.05) and No Field (p<0.05) conditions (**Fig. 3B**). This effect was not identified between exposure conditions at 6-hr (p>0.05) or 24-hr (p>0.05) post-exposure timepoints (**Fig. 3C,D**), indicating that the LTP-EMF produced a transient reorganization of network dynamics rather than long-term shifts in baseline excitability.

**Figure 3.**
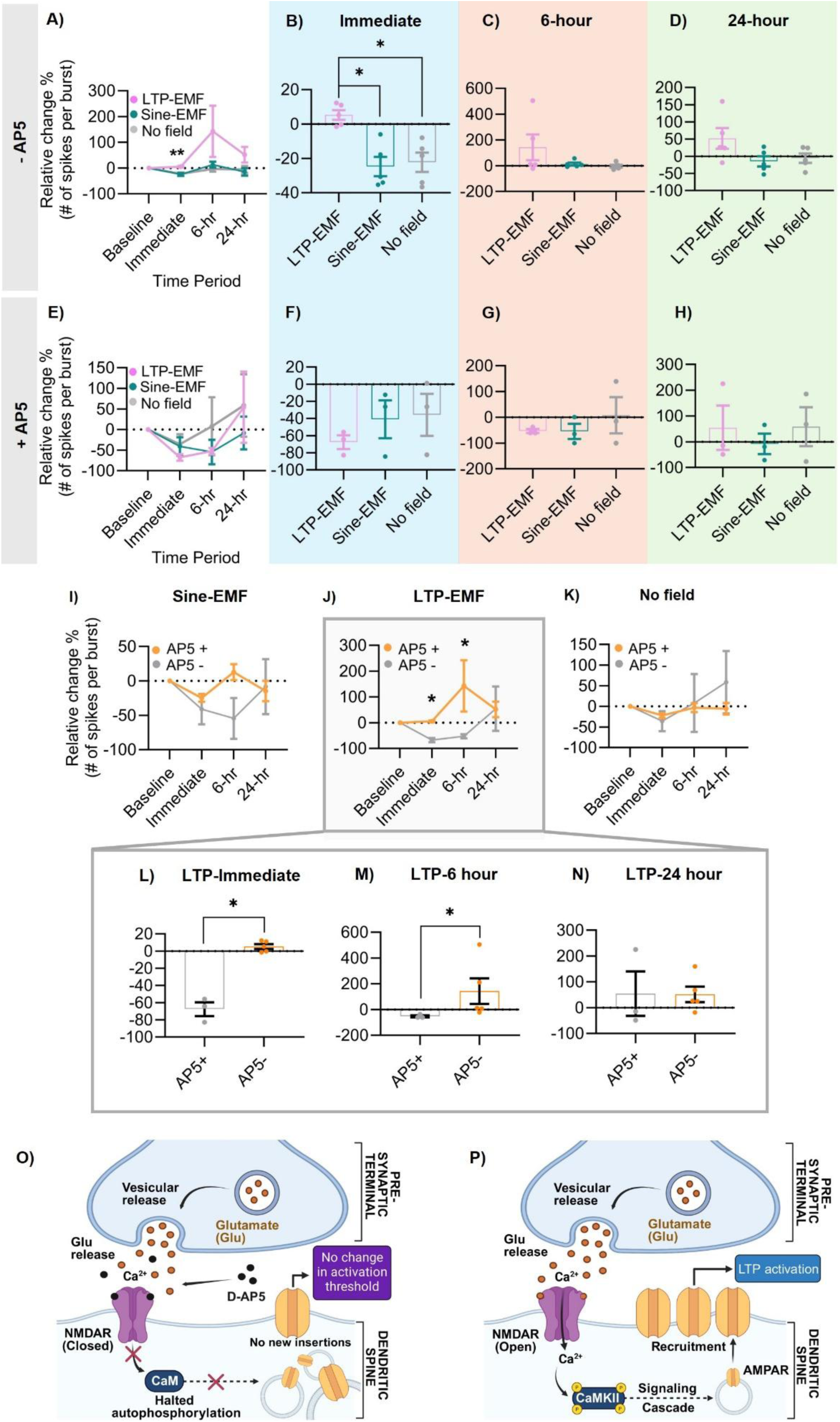
LTP-EMF exposures transiently increased spikes-per-burst with NMDA receptor-dependencies. Primary cortical networks cultured on HD-MEAs were exposed to the LTP-EMF, Sine-EMF, or No Field conditions and electrophysiological recordings were collected to determine spontaneous network properties. (A) A significant field-dependent effect of change in normalized spikes-per-burst was observed across timepoints (n=5/condition). (B) Post-hoc comparisons confirmed a selective increase in spikes-per-burst immediately following LTP-EMF exposure relative to Sine-EMF and No Field conditions. This effect was absent at the (C) 6-hr and (D) 24-hr timepoints. (E-H) In AP5-treated cultures (n=3/condition) (100 µM), no field-dependent effect on spikes-per-burst was detected at any timepoint. (I-K) Within-field comparisons revealed that AP5 selectively suppressed spikes-per-burst in LTP-EMF-exposed cultures (J), with no significant effect in (I) Sine-EMF or (K) No Field conditions. Follow-up analyses within the LTP-EMF condition confirmed AP5-dependent reductions in spikes-per-burst (L) immediately and at (M) 6-hrs post-exposure, with no residual effect at (N) 24-hrs, consistent with the temporal dynamics of LTP early-phase expression. (O) Schematic of the established mechanism underlying AP5-mediated blockade of the LTP-EMF effect, where inhibition of NMDA receptor activation prevents downstream signaling and AMPAR recruitment. (P) Schematic of LTP induction in the absence of AP5. These findings support the hypothesis that the electrophysiological effects induced by the biomimetic LTP-EMF are NMDA receptor-dependent and are attenuated by pharmacological blockade of synaptic plasticity mechanisms. Statistically significant differences are indicated: *p < 0.05, **p < 0.01.

### 2.2 NMDA receptor blockade abolished LTP-EMF-evoked network burst dynamics

If LTP-patterning confers physiological specificity to EMF-based stimulation, the cellular mechanisms underlying its electrophysiological effects should converge with established pathways of activity-dependent synaptic plasticity. NMDA receptor activation is a well-defined molecular requisite for LTP induction, gating the Ca²⁺ influx that triggers downstream signalling cascades and AMPAR trafficking [30]. We therefore asked whether pharmacological antagonism of NMDA receptors would abolish the LTP-EMF-evoked increase in spikes-per-burst. Bath application of the competitive NMDA receptor antagonist AP5 (100 µM) during EMF exposure completely abolished the field-dependent increase in spikes-per-burst, with LTP-EMF cultures showing no significant difference from Sine-EMF or No Field conditions at any timepoint (p> 0.05; **Fig. 3E–H**). Critically, within-field comparisons between AP5-treated and untreated cultures revealed that AP5 selectively reduced spikes-per-burst in the LTP-EMF condition, F(1,6)=11.46, p<0.05 (**Fig. 3J, L**), with no significant AP5-dependent changes in Sine-EMF or No Field cultures (**Fig. 3I, K**). This selectivity began post-stimulation (U=0, p<0.05; **Fig. 3L**) persisted at the 6-hr post-exposure timepoint (U=0, p< 0.05; **Fig. 3M**) and was resolved at 24-hrs (p>0.05; **Fig. 3N**), mirroring the temporal dynamics of early-phase LTP expression.

Together, these data demonstrate that the electrophysiological effects of LTP-patterned EMF exposure are NMDA receptor-dependent and engage excitatory glutamatergic transmission pathways that are mechanistically continuous with those underlying endogenous synaptic potentiation [30] that are known to be attenuated by AP5 (**Fig. 3O,P**).

### 2.3 LTP-EMF exposures transiently decreased network excitability with rebound effects

Increased network excitability is a defining feature of LTP, where responsiveness to input stimulation is amplified or “potentiated” as synaptic contacts strengthen and become more efficient [31–33]. However, before transitioning to a long-term state characterized by reduced activation threshold, neural networks are known to display transient insensitivity to evoked potentials in the immediate phase following stimulation as pre-synaptic vesicles become depleted, receptors desensitize, and ion channel dynamics normalize [34,35]. Having observed the NMDA receptor-dependent increase in spontaneous spikes-per-burst (**Fig. 3**), we predicted that LTP-EMF exposure would be accompanied by a concomitant, transient reduction in the capacity of the network to recruit additional spiking in response to exogenous voltage inputs. To test this, we applied a graded extracellular stimulation protocol (200–800 mV, in 100 mV increments) via MEA electrodes immediately, 6-hrs, and 24-hrs following EMF exposure, and quantified the relative change in active electrode recruitment compared to a 0 mV sham baseline (**Fig. 4A, B**). Here, networks were only ever exposed to 1 stimulation assay; therefore, different chips (n=5/condition/timepoint) were used for each timepoint to eliminate the confound of repeated stimulation.

**Figure 4.**
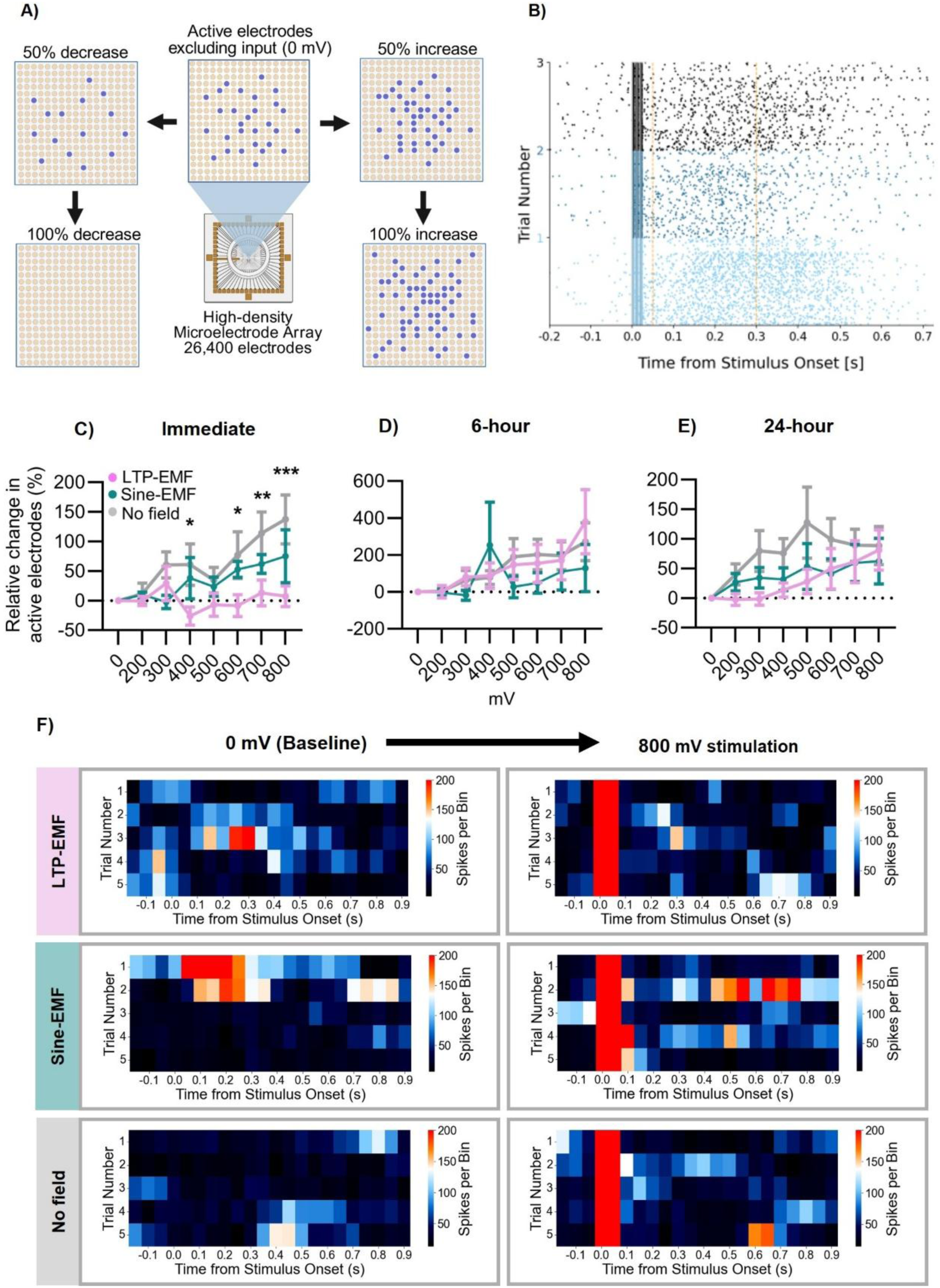
Network responsiveness to voltage inputs was transiently inhibited immediately following LTP-EMF exposures. (A) Schematic depiction of relative changes in active electrode recruitment (± 50-100%) across the HD-MEA array, normalized to the 0 mV sham stimulation condition. (B) A representative raster plot illustrating stimulus-evoked responses; note the evoked spikes after the onset at 0.0 seconds as indicated by the dark vertical lines. (C-E) Relative change in active electrode recruitment (%) as a function of increased stimulation amplitude (0-800 mV) for LTP-EMF, Sine-EMF, and No Field conditions at the (C) immediate, (D) 6-hr, and (E) 24-hr post-exposure timepoints (n=5/condition/timepoint). Values are presented relative to 0 mV sham baseline. Significant field-dependent decreases in active electrodes were observed immediately following LTP-EMF exposure at higher stimulation amplitudes (>400 mV). (F) Representative heat maps of spiking activity contrasting the 0 mV sham (baseline) and the highest input amplitude, 800 mV, illustrating diminished responsiveness to applied voltages among LTP-EMF-exposed networks. Note that LTP-EMF-exposed networks display more spikes per bin at baseline (0 mV) and an insensitivity to input voltages whereas Sine-EMF and No Field controls display relatively inactive baseline states with evoked activity upon stimulation. Statistically significant differences are indicated: *p < 0.05, **p < 0.01, ***p < 0.001.

An analysis of stimulation-dependent changes in active electrodes relative to baseline activity revealed a significant timepoint-by-condition interaction, F(12,72)=6.69, p<0.0001. Immediately following EMF stimulation, neural networks cultured on MEA chips that were exposed to the LTP-EMF pattern displayed a significant decrease in active electrodes relative to controls when stimulated with 400 mV (p<0.05), 600 mV (p<0.05), 700 mV (p<0.01), or 800 mV (p<0.001) (**Fig. 4C**). Interestingly, no voltage-dependent effects could be discerned within the LTP-EMF condition at the immediate timepoint (p>0.05), suggesting an acute insensitivity to stimulation. No significant voltage-dependent changes in active electrodes between EMF conditions were detected at the 6-hr or 24-hr post-EMF exposure timepoints (**Fig. 4D, E**; p>0.05). Stimulation-response plots suggested an elevated level of spontaneous activity during sham stimulation (0 mV) for the LTP-EMF condition, with insensitivity to even the most extreme voltage inputs (800 mV) (**Fig. 4F**). These results demonstrate that LTP-EMF exposures caused an immediate, transient period of insensitivity to applied voltages during the same period that they displayed greater spontaneous spikes-per-burst, suggesting a reversible suppression of network excitability rather than a non-specific or cytotoxic effect.

### 2.4 LTP-EMF exposures promoted delayed transcriptional states consistent with structural remodelling

As the primary cortical networks displayed increased spikes-per-burst and immediate, transient insensitivities to evoked network activity, we next assessed whether these electrophysiological changes were accompanied by transcriptional responses. To characterize the molecular consequences of LTP-EMF exposure, we performed bulk RNA sequencing at two post-exposure timepoints corresponding to established temporal phases of LTP gene expression: 30-min (early response) and 6-hrs (delayed response) [36, 37].

Sample-to-sample Euclidean distance analysis showed that transcriptional profiles clustered primarily by post-exposure timepoint, with less pronounced separation by EMF condition alone (**Fig. 5A**). Differential expression analysis revealed that the strongest field-associated transcriptional response occurred 6-hr after LTP-EMF exposure. Hierarchical clustering of the 50 genes with the lowest adjusted p-values further separated samples by post-exposure time and by EMF condition at 6-hrs (**Fig. 5B**). At 6-hrs, the LTP-EMF versus No Field comparison identified 96 upregulated and 17 downregulated genes (**Fig. 5D**), while the Sine-EMF versus No Field comparison at the same timepoint yielded no significantly differentially regulated genes (**Fig. 5E**). Gene Ontology enrichment analysis of the 96 upregulated genes identified 26 significantly enriched terms (**Supplementary Table S1**). The top enriched processes included regulation of apoptotic signalling pathway, endoplasmic reticulum unfolded protein response (GO:0030968), cellular response to unfolded protein (GO:0034620), cellular response to glucose starvation (GO:0042149), intrinsic apoptotic signalling pathway in response to endoplasmic reticulum stress (GO:0070059), filopodium assembly (GO:0046847), and ER overload response (GO:0006983), driven by genes including *Osgin1*, *Ddit4l*, *Ier3*, *Gdf15*, *Creb3l1*, *Fgf21*, *Hmox1*, *Fosl1*, *Junb*, *Itga5*, *S1pr2*, and *Bdkrb2* (**Fig. 5C**). No pathways were significantly enriched among down-regulated genes. Together, these results suggested that the transcriptional response was waveform-specific and temporally restricted to the delayed post-exposure window.

**Figure 5.**
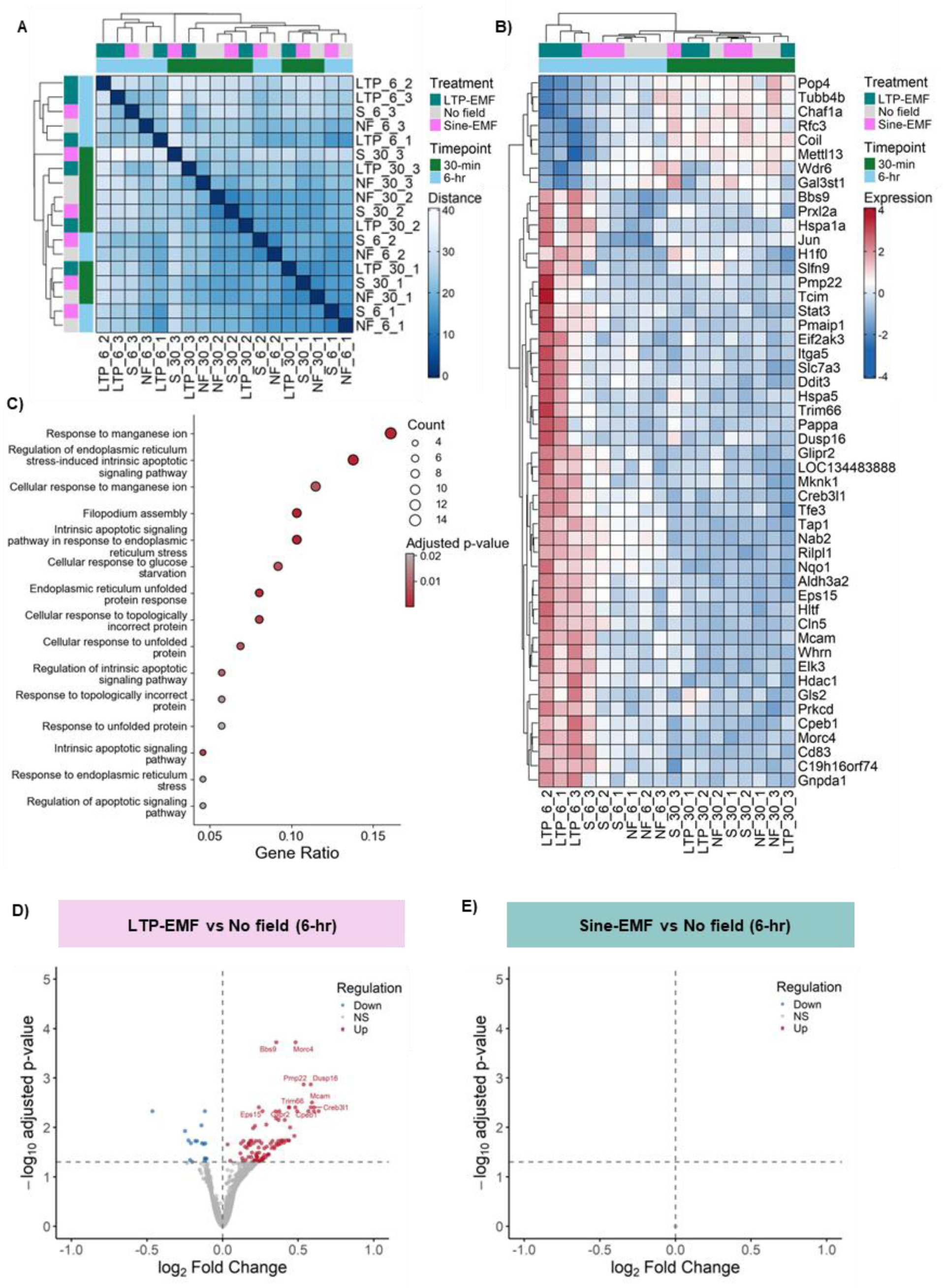
Transcriptomic profiling of primary rat neurons following electromagnetic stimulation. Primary rat cortical neurons were exposed to LTP-EMF (pink), Sine-EMF (cyan), or No Field (grey) for 30-mins, and RNA was collected at 30-mins or 6-hrs post-stimulation for bulk RNA sequencing (n=3/condition). (A) Sample-to-sample Euclidean distance heatmap with hierarchical clustering. Outer colour bars indicate EMF condition; inner bars indicate post-exposure timepoints (green is 30-min; blue is 6-hr). Distance scale ranges from dark blue (minimal distance) to white (maximal distance). (B) Hierarchically clustered heatmap of differentially expressed genes for the LTP-EMF versus No Field comparison at 6-hrs. The 50 genes with the lowest adjusted p-values were used as the basis for sample and gene clustering. The top dendrogram clusters samples by treatment condition and timepoint; the left dendrogram clusters genes by expression profile. Expression values are shown relative to the per-gene mean, with red indicating upregulation and blue indicating downregulation. (C) Gene Ontology pathway enrichment for genes significantly upregulated in LTP-EMF relative to No Field at 6-hrs (top 15 terms shown). Dot size reflects the number of genes contributing to each pathway; x-axis position reflects the gene ratio; colour indicates pathway significance (red = lower adjusted p-value). Volcano plots of differential gene expression for (D) LTP-EMF versus No Field and (E) Sine-EMF versus No Field at 6-hrs. The x-axes show log₂ fold change while the y-axes show −log₁₀ adjusted p-value. Upregulated genes, red; downregulated genes, blue; non-significant genes, grey.

### 2.5 New synapse formation was promoted by LTP-EMF exposures

One of the most conspicuous downstream markers of endogenous LTP is the formation of new synaptic contacts, typically accompanied by coincident spine enlargement and increased receptor density [38]. Together, these structural changes contribute to establishing persistent, long-lasting increased synaptic strength and efficiency [30]. To determine whether LTP-patterned EMF exposure altered synaptic marker organization, we performed immunofluorescence imaging for synaptophysin (SYP), a presynaptic vesicle marker, and postsynaptic density-95 (PSD-95), a scaffolding protein enriched at excitatory post-synaptic sites (**Fig. 6A**). Spatial co-localization of SYP and PSD-95 proteins provides an index of newly formed synapses (synaptogenesis) [39]. We hypothesized that if the temporal signatures embedded in the LTP-patterned EMF stimulation were sufficient to bias plasticity-associated cellular responses, cultures exposed to LTP-EMF would show increased spatial co-localization of SYP and PSD-95 markers relative to Sine-EMF and No Field control conditions.

**Figure 6.**
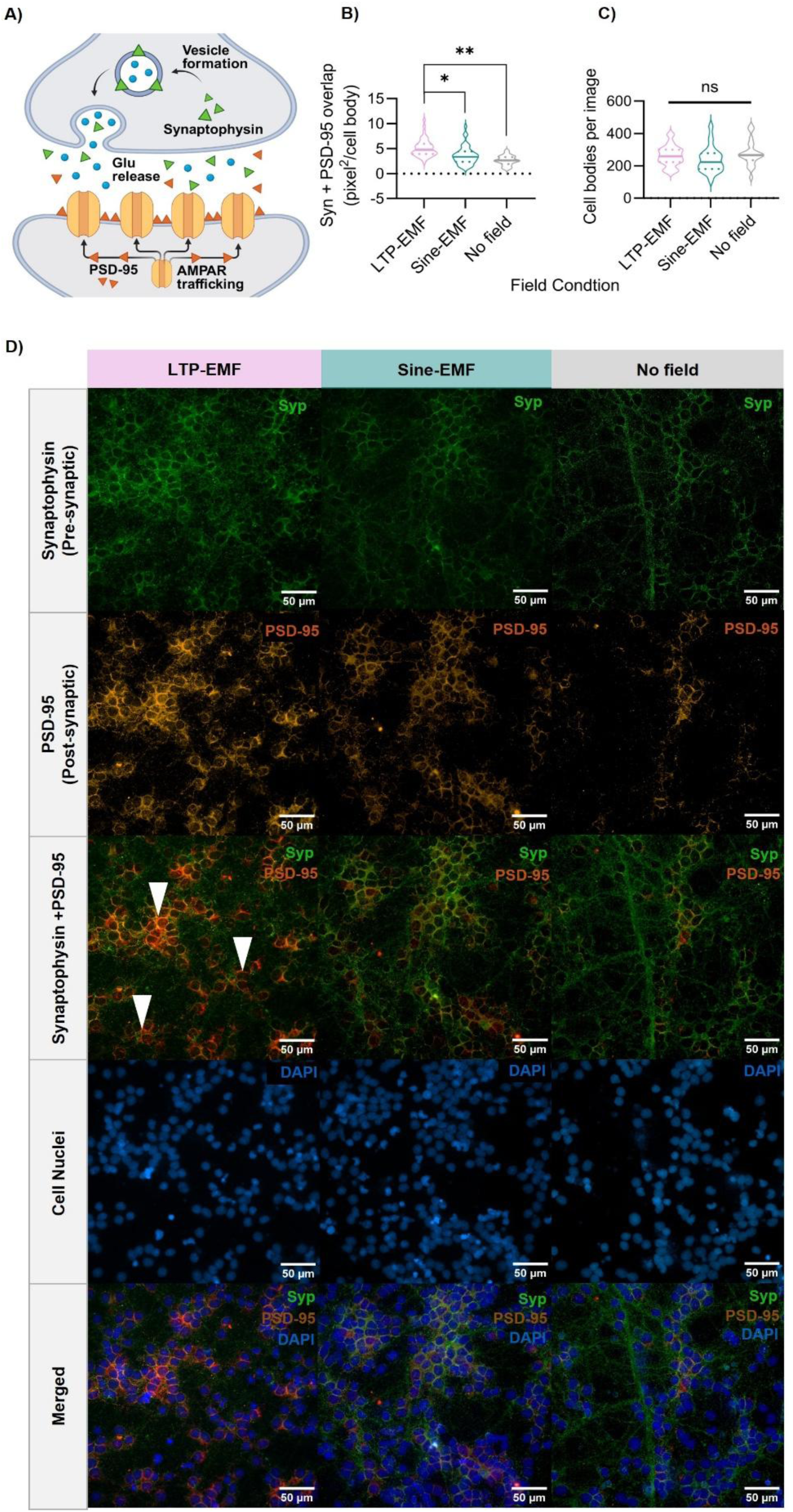
Synaptophysin–PSD-95 colocalization indicative of new synapse formation increased following LTP-EMF exposures. Primary cortical neurons were prepared for immunocytochemistry following 30-min exposure to LTP-EMF, Sine-EMF, or No Field conditions and immunostained for synaptophysin (SYP; green), postsynaptic density-95 (PSD-95; orange), and DAPI (blue). Co-localization of SYP and PSD-95, normalized to DAPI, was quantified as area per cell body (pixels²/cell) from confocal images acquired at 40X. (A) Schematic representation of targeted pre- (SYP) and post-synaptic (PSD-95) markers. (B) Quantification of SYP-PSD-95 co-localization. Post-hoc comparisons confirmed significantly greater co-localization in LTP-EMF cultures relative to Sine-EMF (p < 0.05) and No Field (p < 0.001) controls. Each data point represents the mean of 15 images from one biological replicate (n=3 biological replicates/condition; 45 images per group). (C) DAPI-based nuclear counts confirmed no significant difference in cell density across conditions. (D) Representative immunofluorescence images are provided for all markers with a 50 µm scale bar. White arrows indicate conspicuous clusters of SYP-PSD-95 co-localization in the LTP-EMF condition, where they were quantitatively most prevalent. Statistically significant differences (and null effects) are indicated: ^n.s.^p>0.05,*p< 0.05, **p< 0.001.

After a 30-min exposure to EMF conditions or No Field exposure, monolayers of cortical networks were incubated for 1-hr, fixed, immunostained, and imaged. A one-way ANOVA revealed a significant effect of field condition on co-localized synaptic marker area, F(2,6) = 13.95, p < 0.01 (**Fig. 6B)**. Indeed, LTP-EMF exposure significantly increased cell-body-normalized SYP-PSD-95 colocalization relative to Sine-EMF (p < 0.05) and No Field controls (p < 0.01). DAPI-based cell counts confirmed that cell numbers did not differ between conditions (**Fig. 6C**). These data indicate that LTP-EMF exposure promoted the formation of new synaptic contacts and provide a direct structural correlate for the NMDA receptor-dependent network-level effects observed electrophysiologically.

## Discussion

Low-intensity EMFs, below the millitesla (mT,10^-3^ T) range, are often characterized as insufficient to affect biological systems [40], despite well-known examples of low-intensity biological effects in natural settings, particularly among migratory animals [41]. This disconnect is in part due to a narrow focus on simple waveforms [42] associated with power transmission and electronic infrastructure (e.g., 60Hz) that are distinct from the temporally complex, endogenous EMFs within biological systems. The view that EMFs below the mT range are unlikely to affect biological systems is also related to assumed lower bounds on interaction, based on the strength of Earth’s magnetic field, around 50 µT, and thermal energy (kT) within the cell[43]. Both considerations are intensity-related and call into question the ability for a weak signal to overcome a noisy energetic environment. Of course, the evidence runs contrary to these theoretical limits [41], including demonstrations of µT-strength brain interactions [24, 44]. Perhaps intensity is only one of several relevant parameters that determine biological effects of EMFs. Here, we provide evidence that the temporal structure or pattern complexity of the applied EMF may be a critical parameter associated with µT-strength neuromodulation and other low-intensity bioelectromagnetic interactions.

In the present study, we demonstrated that a µT-strength EMF patterned after neuronal firing signatures associated with LTP produced a distinct response in primary cortical networks, increasing spontaneous spikes-per-burst while transiently reducing recruitment to direct electrical stimulation. This response was not reproduced by a frequency-matched sine wave (100 Hz) and was abolished by NMDA receptor antagonism, suggesting that the effect was not explained by mere field exposure and was dependent on excitatory transduction mechanisms. Rather than producing a non-specific increase in excitability, the LTP-patterned waveform shifted network dynamics in a manner that depended, at least in part, on glutamatergic signaling pathways linked to activity-dependent plasticity.

In a previous study [29], whole-body exposure to a biomimetic LTP-patterned EMF attenuated cortical damage following lithium-pilocarpine-induced seizures in rats. The authors proposed that field-sensitive disruptions of Ca^2+^ binding, upstream of excitotoxic apoptotic and necrotic pathways, could account for enhanced cell survival, citing earlier work [45] showing that pulsed µT-strength EMFs altered calmodulin-dependent myosin phosphorylation. In the present study, we identified NMDA receptors, which are predominantly Ca^2+^ permeable ion channels, as a contingent factor in the LTP-EMF effect. Consistent with early reports of memory disruption in rodents exposed to LTP-patterned EMFs [27, 28] and the anti-excitotoxic properties of the same pattern in a rodent model of epilepsy [29], as well as previous demonstrations that µT-strength EMFs can disrupt electrical activity in hippocampal slice cultures [46], we agree that calcium influx is likely a critical physiological bottleneck for the effects of the LTP-EMF exposures. A similar conclusion is supported by longstanding evidence of calcium-dependent effects associated with time-varying, µT-strength EMFs in other cell types [47–49].

This may provide a cellular explanation for earlier behavioural studies in which LTP-patterned EMFs disrupted contextual fear memory in rodents [27, 28]. Because these fields were delivered during temporal windows relevant to learning and memory formation, their effects may depend not only on the intrinsic structure of the waveform, but also on the timing of its application relative to ongoing neurophysiological processes. Memory encoding and early consolidation are highly time-dependent, NMDA receptor-sensitive processes that require coordinated excitatory drive, Ca²⁺ influx, and synaptic plasticity. The *in vitro* results reported here support the hypothesis that the memory-disrupting mechanism observed in animals exposed to LTP-patterned EMFs is likely an interference of excitatory, NDMA receptor-dependent mechanisms that drive time-sensitive memory encoding processes. Thus, biomimetic EMFs may exert their strongest effects when their temporal structure intersects with biological windows in which endogenous plasticity mechanisms are actively recruited.

The observed dissociation between spontaneous (**Fig. 3**) and evoked (**Fig. 4**) was noteworthy and suggestive of an effect tied to excitability. Increased spikes-per-burst might be interpreted as increased excitability or increased recruitment if considered in isolation – either the same cells are generating more spikes within a burst or additional cells are joining the network-level response. However, the simultaneous reduction in active electrode recruitment to applied voltages suggests a more specific, state-dependent effect tied to excitability. One interpretation is that LTP-EMF exposure engages mechanisms at the cellular scale that, in turn, affect network dynamics. The period of insensitivity that we observed at the immediate time point, within 45-60 minutes of initiating the EMF exposure and following post-exposure activity and network assays, is within the temporal range of previously reported refractory periods for plasticity in LTP-induction paradigms [50]. If LTP-EMF exposures diminish excitability, one expected effect would be the interference of memory encoding processes as was previously observed in similarly exposed rodents [27, 28]. If the LTP-EMF pattern is transiently blocking excitability, future applications of the protocol may be timed to engage distinct stages of memory processing, including encoding, consolidation, and retrieval. This does not establish the precise mechanism of coupling, but it is consistent with a reversible reorganization of functional network state that may lead to downstream structural modifications.

The molecular and imaging data provide converging support for this interpretation. Indeed, the transcriptional response was strongest at the 6-hr timepoint and was enriched for pathways associated with adaptive cellular remodeling, including unfolded protein response, endoplasmic reticulum stress, metabolic regulation, redox-related processes, and filopodium assembly. The data suggest that LTP-patterned EMF exposure induced a delayed cellular response compatible with structural remodeling that followed the immediate electrophysiological perturbations within the network. Similarly, increased synaptophysin-PSD-95 co-localization following the LTP-EMF exposure provides a microstructural correlate of synaptic plasticity – predictable downstream consequences of excitatory modulation and transcriptional shifts toward structural remodeling. Together, the observed effects following LTP-EMF exposure suggest modulated excitability with downstream synaptic plasticity.

Several considerations should guide interpretation of the present results. Primary cortical cultures provide a tractable reductionist model for identifying cellular responses, but they do not reproduce the anatomical organization, cell-type diversity, long-range connectivity, or behavioral context of intact neural circuits. The Sine-EMF condition served as a frequency-matched comparator, but it was not matched to the LTP-EMF for duty cycle, pulse timing, or polarity structure. The present study therefore identifies waveform-condition-dependent effects but does not isolate which individual feature of the complex waveform is necessary or sufficient. Future studies using scrambled LTP patterns, duty-cycle-matched pulse trains, polarity-matched controls, amplitude-matched fields, and state-dependent stimulation paradigms will be needed to define the relevant waveform parameters.

The central implication of these findings is that the parameters that determine EMF-based neuromodulation may not be fully explained by field intensity, frequency, or exposure duration alone. Although, the LTP-EMF and Sine-EMF conditions shared a 100 Hz component, only the biomimetically patterned LTP-EMF produced distinct electrophysiological, transcriptional, and microstructural outcomes, while a µT-strength sine EMF was largely comparable to the No Field condition. This suggests that features such as pulse timing, interstimulus intervals, duty cycle, and polarity structure may influence how weak EMF inputs interact with neural networks and other biological systems. In this context, complex waveforms may contain biologically relevant, information-rich temporal structure that interact with the electrochemical dynamics of excitable tissue. This view is consistent with evidence that weak endogenous-scale fields can bias spike timing and entrain network activity [17], and converges with studies showing that low-intensity, environmental EMFs can modulate brain activity [51].More broadly, they support a framework in which biomimetic EMFs are treated as temporally structured biophysical signals whose effects depend not only on the pattern delivered, but also on when that pattern intersects with endogenous windows of plasticity. Defining this relationship between waveform structure and physiological timing may be essential for designing more targeted, non-invasive strategies to modulate neural activity.

## Experimental Methods

### Primary cortical neuron isolation and culture

Primary cortical cells were isolated from embryonic (Day E18) Sprague-Dawley rat brains. All procedures involving animals were conducted following the guidelines established by the Canadian Council on Animal Care (CCAC) and were approved by Wilfrid Laurier University’s Animal Care Committee. Timed-pregnant rats were euthanized by way of external cervical dislocation, and embryos were rapidly extracted under sterile conditions. Embryos were transferred to a sterile Petri dish containing ice-cold, calcium- and magnesium-free Hank’s Balanced Salt Solution (HBSS) (Gibco™, Cat# 14175095). Brains were carefully dissected under a stereomicroscope, and cortical tissue was isolated by removing the meninges and separating the cortical hemispheres from the underlying subcortical structures. Cortices from multiple embryos were pooled in fresh, ice-cold HBSS for enzymatic dissociation. The pooled cortical tissue was incubated in 0.25% trypsin at 37 °C for 20 minutes. Following digestion, the tissue was treated with 10% fetal bovine serum (FBS) in 1X DPBS (Gibco, Cat# 14190144) to deactivate digestion. Mechanical dissociation was performed by gentle trituration using serological pipettes to obtain a single-cell suspension. Cells were pelleted by centrifugation at 200 × g for 5 minutes and resuspended in Neurobasal™ medium (Gibco, Cat# 21103049) supplemented with 2% B-27™ (Gibco, Cat# 17504044), 0.5 mM GlutaMAX™ (Gibco, Cat# 35050061), and 1% penicillin-streptomycin (Gibco, Cat# 15140122). Cell suspensions were then passed through a 40 μm nylon mesh filter to remove tissue debris and undissociated aggregates. Viable cells were counted using the Trypan Blue exclusion method. Following dissociation and counting, cells were resuspended in plating medium at two cell concentrations: 3 × 10⁶ cells/mL for MEA experiments and 10⁶ cells/mL for RNA sequencing, and immunofluorescence. Assay-dependent culture specifications are detailed elsewhere, wherever relevant.

### Electromagnetic Stimulation Exposure Chamber

Applied EMFs were delivered within an exposure chamber that remained inside of an incubator. The exposure chamber was a 5.2” x 5.2” x 5.2” acrylic box, assembled with 4 nylon M3 x 8mm bolts for legs. Hinges for the acrylic lid were fastened with adhesive. Each of the independent channels (1-3) were wired with two coils in series, such that the magnetic poles were aligned. The exposure chamber box was wired through an RJ45 connector and a mini-B 5-pin to a digital-analog converter (DAC) that was responsible for converting customized wave patterns into EMFs. EMF patterns were designed using Windows digital analog converter software (WinDAC, custom software). These patterns were expressed as digital values ranging from 0-255 over time (ms). The digital values represented a voltage range of (±5V). Therefore, one digital value was equivalent to ∼40 mV. Patterns created in this software were then uploaded to a DAC, generating current to pairs of wrapped solenoids surrounding the exposure chamber. Each bobbin of a solenoid was 3D printed in PLA plastic. After sanding/smoothing the bobbin, it was wound with 3550 turns of Elektrisola P155 magnet wire (40 AWG/0.084mm diameter) on a custom coil-winding machine. Adhesive was applied to the finished windings to keep them in place while the coil was completed. After winding, a 2.54 mm header connector was fixed to the coil body with adhesive. The magnet wire was then wound around each leg of the connector and soldered. The core of each coil was an M3 x 25mm bolt and nut, which together brought the inductance of the coil to 155 mH. Each coil was then placed inside a protective plastic casing for attachment to the exposure chamber.

### Electromagnetic Stimulation Patterns

Two waveforms were used as EMF patterns: LTP and Sine (100 Hz). The LTP-EMF pattern was a normalized equivalent of neural firing recorded during learning tasks in rats [52, 53] and has been previously described as firing patterns based on two physiological features of the hippocampus, including complex spike discharges and theta rhythms [19, 54]. This pattern composition involves a primer stimulus followed by a 150 ms interstimulus interval, followed in turn by four cycles of a 100 Hz stimulation. This sequence combines both low- and high-frequency strategies for inducing LTP as a primer pulse followed by burst patterns [55, 56]. The duration of each discrete stimulus in the series of pulses was 5 ms and, coded as digital values 128-255 (0V - 5V of output). The total duration of the pattern occurred over 225 ms; thus, it could be pulsed twice by each pair of solenoids over a 450 ms period. The Sine-EMF pattern was a continuous 100 Hz sine wave cycling between digital values 0-255 (±5V range) and lasting the same duration as the LTP pattern.

Pairs of solenoids positioned around the EMF chamber box pulsed sequentially in each of the three spatial planes: X, Y, and Z for 450 ms each (Pairs 1A-B, 2A-B, 3A-B, see **Fig. 2**). This was followed by a combined pulse of all three planes for an additional 450 ms. The total duration of the full stimulation cycle (Pair 1, then 2, then 3, then 1+2+3) was 1.8 seconds. Computing digital values from 0-127 induced a negative flow, where the north poles were in the direction of solenoid B. Alternatively, digital values coded from 128-255 had their poles reversed under positive voltage. These directional changes created fundamental distinctions between the field conditions used in this study. For instance, the Sine-EMF wave alternated polarity every 5 milliseconds, completing a full cycle every 10 milliseconds. This resulted in a continuous and symmetrical oscillation between positive and negative phases throughout the entire stimulation period (**Fig. 2**, bottom left panel). In contrast, the biomimetic LTP-EMF was more complex. Rather than alternating polarity, it fluctuated between positive current flow and periods of no current (i.e., off states) as a monophasic stimulus (**Fig. 2**, bottom right panel). Importantly, the direction of current in the LTP-EMF was unidirectional; however, at specific time points – outlined in the waveform design – this current was interrupted rather than reversed. Notably, control plates or chips (No Field condition) were placed in the exposure chamber for the same durations without turning on signal generators to mimic all environmental conditions except EMF exposure.

A DC/AC 3-axis milligauss meter (AlphaLab Inc.) was used to validate the magnetic field strength within the exposure chamber and confirm the presence of weak electromagnetic flux. During application of the LTP-EMF, the measured field intensity between two solenoids of a single-axis reading fluctuated between 1273 and 1835 milligauss (mG), corresponding to approximately 127.33 to 183.5 μT. In contrast, the 100 Hz Sine wave condition produced a steady field intensity of 173.0 μT. The difference in intensity between the two field patterns, integrated over the same period of time, was consistent with their waveform characteristics, holding all other parameters constant. While the Sine-EMF maintained a continuous 100 Hz periodicity throughout the exposure, the LTP-EMF waveform delivered bursts of rapid change interspersed with periods of non-stimulation (interstimulus intervals mimicking features of biological signaling), resulting in a lower overall time-averaged field strength. With the options to hold frequency or time-averaged field strength constant as a non-complex EMF comparator, the 100 Hz Sine pattern was selected as a simplified, frequency-matched version of the LTP stimulus without biologically relevant pulsations or interstimulus delays.

### High-Density Microelectrode Array (HD-MEA) Electrophysiology

HD-MEA experiments were performed with assay-specific coating, seeding, and culturing conditions. Prior to cell seeding, each HD-MEA chip (MaxOne, MaxWell Biosystems Inc.), along with its corresponding holder and lid, was removed from sterile packaging and submerged in 70% ethanol for 30 minutes to ensure sterilization. Following ethanol treatment, all components were rinsed three times with deionized (DI) water to remove any residual ethanol. The holders and lids were then air-dried completely inside a biosafety cabinet under sterile conditions. Once dry, the chips were immersed in a 1% Tergazyme^TM^ (Alconyx^TM^, Cat# 04-322-11C) solution prepared in DI water and incubated for two hours. After this step, the chips were rinsed three times with DI water, then submerged again in 70% ethanol, followed by three additional DI water rinses. Finally, the holders and lids were air-dried once more inside a biosafety cabinet under sterile conditions. Sterilized chips were transferred to 140-mm Petri dishes, each containing a smaller 35 mm Petri dish filled with DI water positioned at the center to serve as a humidity chamber, thereby minimizing evaporation of culture medium during incubation. Each chip was preconditioned by filling it with 600 µL of Neurobasal formulation (described elsewhere). Chips were incubated for 24 hours at 37°C, 5% CO₂, and 95% relative humidity to allow the surface to equilibrate and improve cell attachment conditions. Following pre-conditioning, the medium was aspirated, and each chip was rinsed once with DI water and allowed to dry. Then, 50 µL of a 0.5 mg/mL poly-D-lysine (PDL) solution (Sigma Aldrich, Cat# A-003-M) prepared in 1X Dulbecco’s Phosphate-Buffered Saline (DPBS) was applied to the central electrode area of each chip to promote cellular adhesion. Chips were incubated for an additional 24 hours. After incubation, the PDL solution was aspirated, and chips were washed several times with DI water to remove excess coating reagent. Chips were then allowed to dry at room temperature for 1 hour before cell seeding.

Following chip preparation, 50 µL of the cell suspension from primary neuron isolation procedure (described elsewhere) – containing approximately 150,000 neurons – was carefully pipetted onto the center of the HD-MEA surface. Chips were returned to the incubator for 1 hour at 37°C, 5% CO₂, and 95% humidity to allow cells to adhere to the electrode array. Subsequently, an additional 550 µL of plating medium was added to each chip, bringing the total volume to 600 µL per chip. On the first day *in vitro* (DIV) post-seeding (DIV 1), the medium was fully replaced with a serum-free maintenance medium consisting of BrainPhys™ Neuronal Medium (STEMCELL Technologies, Cat# 05790), supplemented with 2% NeuroCult™ SM1 (STEMCELL Technologies, Cat# 05711), 0.5 mM GlutaMAX™, 15 mM D-(+)-glucose (Thermo Fisher, Cat# J60067-AK), and 1% penicillin-streptomycin. Partial media replacements were performed every three days until DIV 12 to maintain cultures.

At DIV 12, neuronal activity was recorded from all prepared chips to assess their suitability for downstream experimentation. Since post-treatment outcomes were to be evaluated relative to each chip’s baseline, the primary inclusion criterion was the presence of bursting activity, indicating a functionally connected network. To begin, chips were carefully removed from their 140-mm Petri dish humidity chambers and placed onto the MaxOne HD-MEA recording platform, which was maintained inside an incubator at 37°C, 5% CO₂, and 95% relative humidity. Within the MaxWell Biosystems MaxLab Live software, each chip was initialized, offset for stabilization, and a high-pass filter was applied to exclude signals below 300 Hz. Recorded activity was considered biologically relevant if two criteria were met: 1) the offset value remained between 0 and ±5 arbitrary units within the software, and 2) the baseline noise level was maintained at 4 ± 2 µV root mean square (rms). Once calibration was complete, the raster plot was used to visualize real-time spiking across the array. Chips that exhibited spontaneous burst activity, rather than isolated spikes, were deemed suitable for experimental procedures. Based on this criterion, chips displaying robust bursting were selected and randomly assigned to experimental conditions.

HD-MEA recordings began with baseline electrophysiological assessment before EMF exposure. First, a ten-minute activity scan was performed using a checkerboard configuration, which activates every other electrode across the array (13,200 electrodes). This scan enabled detection of spontaneous spiking across a broad spatial range without overloading the readout channels. Immediately following the activity scan, a five-minute network assay was conducted. This assay was only performed on electrodes identified as active in the preceding scan and captured high resolution information on burst patterns to determine numbers of spikes contained within each network burst. The spikes-per-burst metric was treated as a marker of recruitment. Upon completion of baseline recordings, each chip was exposed to its designated EMF condition (Sine-EMF, LTP-EMF, No Field) for 30 minutes inside the incubator (maintained at 37°C, 5% CO₂, and 95% humidity). After the EMF exposure period, each chip was promptly returned to the MEA device. A 2-minute calibration period was allowed to stabilize the signal and permit cellular adaptation to baseline conditions. Subsequently, the activity scan and network assays were repeated using the same configurations as during baseline to allow direct pre-post comparisons.

Following the baseline activity recordings, a stimulation assay was performed to evaluate network excitability. Regions of interest were identified based on firing rate data collected during the post-exposure spontaneous activity scan. These regions were then subjected to extracellular electrical stimulation using a graded voltage protocol ranging from 200 to 800 mV, in 100 mV increments, with 0 mV serving as the control condition. Stimulation parameters followed the default configuration provided by the MaxOne platform: five bursts per stimulation train, each burst consisting of three pulses spaced 10 ms apart, with a 1-second interval between bursts. The order of the graded voltage stimulations was randomized. For each stimulation voltage, neural activity was recorded for 1 minute before and after stimulation to allow normalization of the evoked response relative to baseline firing at 0 mV.

All post-exposure assays – including the activity scan, network assay, and stimulation assay – were performed immediately and repeated 6-hrs and 24-hrs after field exposure to assess the persistence or reversibility of any EMF-induced effects. The “immediate” timepoint is defined as within 2 minutes of stopping the field (or sham) exposure, which includes the time to transport the MEA chip from an exposure incubator to a measurement incubator within the same room and initiate the first assay. No additional field exposures were applied at later timepoints; the recordings were performed under standard incubation conditions to enable longitudinal comparison.

### Blockade of NMDARs with AP5

A subset of MEA electrophysiology experiments involved the use of chemical blockers to attenuate EMF-induced responses. Following standard procedures up to DIV 12, each chip then underwent a baseline activity scan, followed by a network assay, once deemed suitable for experimentation based on the presence of spontaneous bursting. To evaluate whether EMF-induced activity could be blocked via NMDA receptor antagonism, a 1 mM solution of (AP5) (Sigma-Aldrich, Cat# 165304) was diluted to 100 μM directly into the culture medium of the experimental chip. The chip was then incubated at 37°C, 5% CO₂, and 95% relative humidity for a 15-minute period. Immediately following drug incubation, the chip was exposed to its designated field condition – either a 100 Hz sine wave (Sine-EMF), LTP-patterned field (LTP-EMF), or a No Field control – for 30-mins under standard incubation conditions. After field exposure, the chip was promptly transferred back to the MEA platform. A 2-min calibration period was allowed to stabilize the signal and enable cells to adapt to baseline conditions. The activity scan, network assay, and stimulation assay previously described were then repeated to assess any immediate post-treatment changes in network activity and excitability under AP5 treatment. To eliminate any residual drug effects, a full medium exchange with fresh BrainPhys formulation was performed following the post-treatment recordings. A final set of recordings – including the activity scan, network assay, and stimulation assay – was repeated 6-hrs and 24-hrs later to assess the persistence or recovery of network behaviour once the blocking agent had been washed out.

### RNA Sequencing

RNA sequencing was performed to assess the transcriptomic profiles of cells exposed to the EMF conditions. Beginning with cell seeding, 200 µL of the respective suspension – containing approximately 200 thousand neurons – was pipetted into inner and outer regions of pre-coated 48-well plates to sample multiple regions of the EMF-exposure chamber (Corning, Cat# 356509). The outer wells included: A2, A7, B1, B8, E1, E8, F2, F7. The inner wells included: C3-C6 and F3-F6. On DIV 1, the medium was fully replaced with fresh Neurobasal formulation, followed by partial media replacement every three days until DIV 7.

After DIV 7, each of the four plates was subjected to its designated EMF condition for 30 minutes, either the LTP-EMF, Sine-EMF, or a No Field control. Following stimulation, one plate from each condition was processed for immediate RNA extraction, while the remaining plates were returned to the incubator for six additional hours before RNA collection to assess delayed gene expression responses. At both time points, cells were gently washed once with 1X phosphate-buffered saline (PBS) (Gibco™, Cat#10010023) to remove residual media. Total RNA was then extracted using the Qiagen RNeasy Mini Kit according to the manufacturer’s protocol. Wells were pooled corresponding to their outer/inner region. RNA concentration and purity were assessed using a Nanodrop spectrophotometer, and only samples with A260/A280 purity values within ±0.10 of 2.00 were deemed acceptable for downstream analyses.

The genome of *R. norvegicus* was indexed via the STAR aligner [57], maintaining splice junction information. Genome indexing was performed based on a read length of 101 bp, corresponding to the longest read length in the trimmed dataset. Sample quality and duplication rates were assessed with FastQC. Reads were aligned to the indexed genome using the STAR aligner, and alignment metrics were summarized with MultiQC [58]. Gene-level counts were generated using the featureCounts function from the Rsubread package [59].

Count matrices were imported into R and processed with DESeq2 [60]. Size-factor normalization was applied to account for sequencing depth and RNA composition, followed by the variance stabilizing transformation (VST) function to convert the data from heteroscedastic to homoscedastic. VST-transformed counts were used for exploratory analyses, including hierarchical clustering based on Euclidean distances, visualized with a sample-distance heatmap.

Treatment conditions were defined by magnetic field exposure (No Field, Sine-EMF, or LTP-EMF) and by time post-exposure (30-min or 6-hr). Differential expression contrasts compared each field condition to its time-matched No Field control, and within each field condition, 30-min was contrasted to 6-hr. For each contrast, differential expression results were extracted from the DESeq2 object [60].

Log2-fold changes and p-values were calculated for all genes, and p-values were adjusted for false-discovery rate using the Benjamini-Hochberg method [60, 61]. Genes with adjusted p-values < 0.05 were considered significantly differentially expressed and were classified as either upregulated or downregulated. When more than five genes were upregulated or downregulated, pathway enrichment was performed using the enrichGO function from the clusterProfiler package, applying thresholds of p<0.05 and q< 0.2 [62]. A hierarchical heatmap was generated using all differentially expressed genes for each contrast to visualize sample clustering across treatment groups.

### Immunocytochemistry and Flourescence Imaging

Immunocytochemistry was performed to confirm the presence of increased synapses associated with EMF conditions expected to induce LTP. Beginning with cell seeding, 200 µL of the single cell suspension – containing approximately 200 thousand neurons – was pipetted into glass-bottom chamber coverslips (Ibidi, Cat# 80826). On DIV 1, the medium was fully replaced with fresh Neurobasal formulation, followed by partial media replacement every three days until DIV 7. Before seeding, coverslips were coated with 0.1 mg/mL PDL solution and incubated for 24 hours (37°C, 5% CO₂, 95% humidity). After incubation, the PDL solution was aspirated, and the plates were washed several times with DI water to remove excess coating reagent.

On DIV7, neurons were exposed to their respective EMF conditions for 30 minutes using a custom-designed stimulation chamber. Immediately following stimulation, all cultures were returned to standard incubation conditions (37°C, 5% CO₂, 95% humidity) and maintained for an additional hour to allow for early stages of synaptic protein expression. This delay accounts for the temporal dynamics of structural plasticity, as proteins such as synaptophysin and PSD-95 [63]. After incubation, the media was aspirated, and cells were gently washed once with 1X PBS. Neurons were then fixed with 4% paraformaldehyde (PFA) (Thermo Fisher, Cat# J19943.K2) for 10 minutes at room temperature, followed by three washes with 1X PBS, each conducted on an orbital shaker for 5 minutes. Fixed cells were permeabilized and blocked by incubation in blocking buffer consisting of 1% bovine serum albumin (BSA) (Sigma-Aldrich, A4737), 2% goat serum (Gibco, Cat# 16210064), and 0.2% Triton X-100 (Sigma-Aldrich, Cat# X100) in 1X PBS for 1 hour at room temperature.

After blocking, cells were incubated overnight at 4°C with primary antibodies diluted in blocking buffer. The following antibodies and concentrations were used: Synaptophysin: rabbit anti-synaptophysin (Abcam, Cat# ab32127), 0.1 µg/mL and PSD-95: mouse anti-PSD-95 (Abcam, Cat# ab192757), 1 µg/mL. On the following day, the primary antibody solution was aspirated, and each well was washed three times with 1X PBS. Corresponding secondary antibodies, including Goat anti-mouse IgG H&L (Alexa Fluor® 594), preadsorbed (Abcam, Cat# ab150120) for PSD-95, and Goat anti-rabbit IgG H&L (Alexa Fluor® 488) for synaptophysin, diluted 1:1000 in blocking buffer, were then applied under low-light conditions and incubated overnight at 4°C. Following secondary incubation, wells were washed three additional times with 1X PBS. DAPI (4′,6-diamidino-2-phenylindole) was diluted to 1 µg/mL and applied to each sample for 5 minutes to visualize nuclear staining. DAPI solution was then removed, and cells were washed three times with 1X PBS. Finally, samples were stored in 1X PBS at 4°C and protected from light until imaging.

Fluorescence imaging was performed using a Nikon Eclipse Ti2 inverted microscope equipped with NIS Elements software. Multichannel images were acquired using standard fluorescence filters for FITC (Alexa Fluor 488), TRITC (Alexa Fluor 594), and DAPI. All images were captured using a 40x objective, with a uniform exposure time of 300 ms per channel and laser intensity set at 15%. For each well, three representative fields of view were selected, focusing on areas with uniform and healthy cell distribution. Imaging was conducted across six wells per coverslip, totalling 15 images per biological replicate, of which there were 3 total. Images were saved as ND2 files for further analysis.

Quantitative image analysis was conducted using ImageJ 1.54p (Java 21.0.7; 64-bit) (National Institutes of Health). Raw ND2 files were imported and separated into individual fluorescence channels, producing grayscale images corresponding to synaptophysin (488 nm), PSD-95 (594 nm), and the nuclear marker DAPI. To assess colocalization between synaptic markers, grayscale images for synaptophysin and PSD-95 were independently converted into binary masks using a standardized intensity threshold of 3.5%, allowing consistent detection of signal across samples. A third binary image was generated using the Image Calculator function to identify pixels in which both synaptic signals overlapped, providing a measure of potential synaptic contact zones. The area of colocalization was quantified in pixels², with spatial resolution calibrated such that one micron equaled 6.17 pixels based on the microscope’s optical specifications. For cell density estimation, the DAPI channel was also converted to a binary image to isolate nuclear profiles. A watershed segmentation algorithm was applied to separate closely apposed or overlapping nuclei, ensuring each cell body was counted as a discrete particle. Subsequent particle analysis was performed with size parameters set between 10–500 µm² and a circularity range of 0.3 to 1.0, where higher circularity values indicated more spherical nuclei.

### Data processing and statistical analyses

Unless specified elsewhere, all statistical analyses were performed in R v4.6 and GraphPad Prism v11.0.1. Data are presented as mean ± standard error of the mean (SEM). Data were first assessed for normality using the Shapiro-Wilk test. In cases where data deviated from normality, non-parametric statistical tests were performed. For comparisons involving more than two groups, one-way ANOVA with appropriate Tukey’s post-hoc multiple comparison testing was used for normally distributed datasets, whereas Kruskal-Wallis tests with Welch’s correction followed by Dunn’s multiple-comparisons were used for non-normally distributed data. Repeated measurements collected across time points or stimulation intensities were analyzed using repeated measures ANOVA. For transcriptomic analyses, differential gene expression was assessed using the DESeq2 package, with p-values adjusted for multiple testing using the Benjamin-Hochberg false discovery rate correction.

## Data Availability

All data needed to evaluate the manuscript are present in the manuscript or the supplementary materials.

## Acknowledgements

The authors acknowledge technical contributions and helpful insights from John Carscallen, Mattia Bonzanni, and Bruce McKay. The research was supported by grants from the Natural Sciences and Engineering Council of Canada (NSERC) including Discovery Grant RGPIN-2021-03783 (to NJM) and RGPIN-2022-04162 (to NR). NR also received support from the New Frontiers in Research Fund – Exploration program (NFRFE-2023-00568). NR and NJM acknowledge support from the Allen Discovery Center at Tufts University. NJM acknowledges support from the Canada Research Chairs program.

## Author Contributions

CK, JSJ, and NVO performed experiments. CK and JSJ equally contributed to data analysis. CK, NR, and NJM prepared the figures. NJM and NR wrote the manuscript. All authors approved the final manuscript. NR and NJM contributed equally to the project.

## Supplemental Tables

**Table 1.**
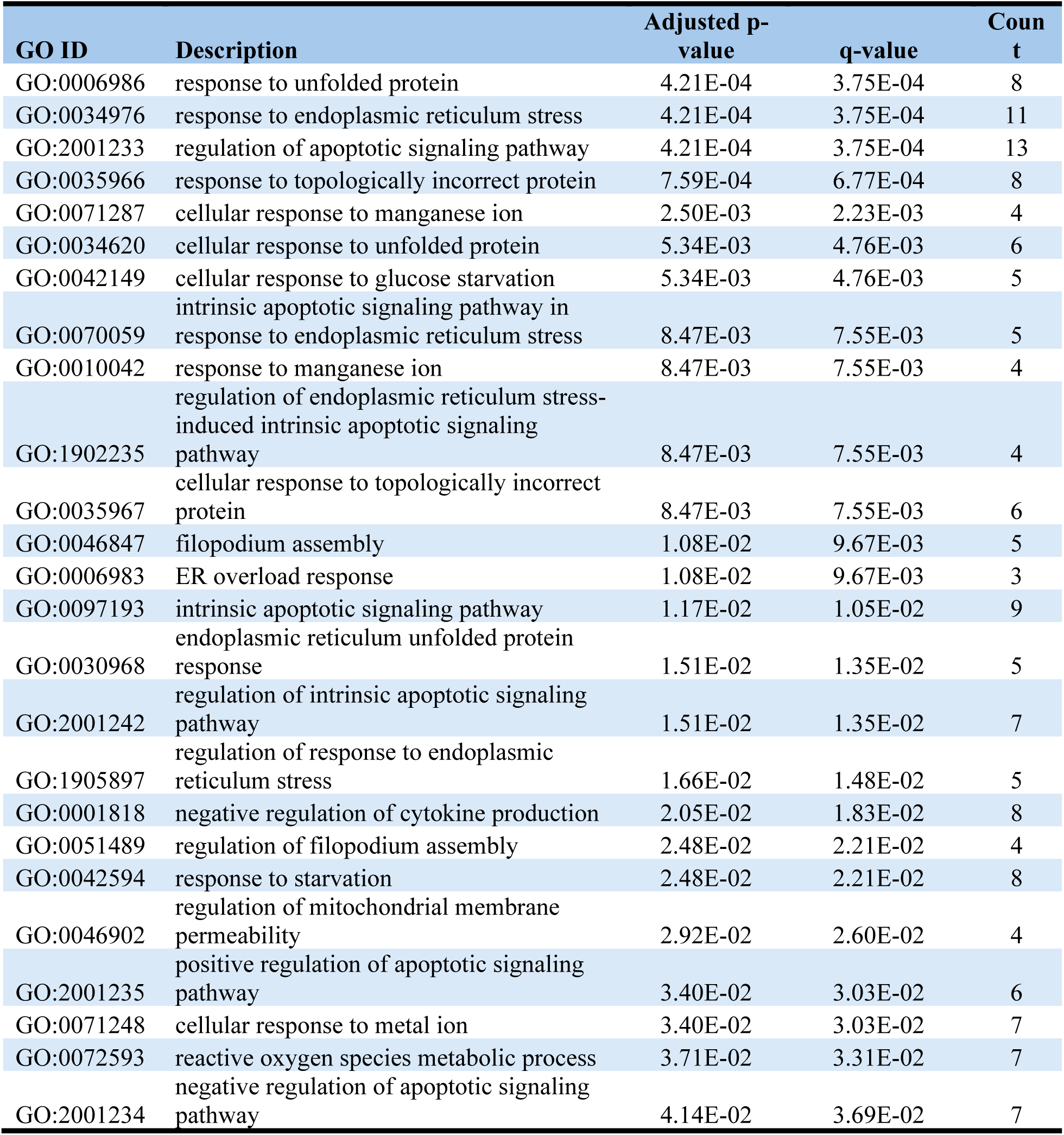
GO pathways significantly enriched in the upregulated genes found in the comparison between LTP-EMF and No Field at 6 h.

